# Helical Ordering of Envelope Associated Proteins and Glycoproteins in Respiratory Syncytial Virus Filamentous Virions

**DOI:** 10.1101/2021.08.04.455049

**Authors:** Michaela J. Conley, Judith M. Short, Joshua Hutchings, Andrew M. Burns, James Streetley, Saskia E. Bakker, Hussain Jaffery, Murray Stewart, B. Joanne Power, Giulia Zanetti, Rachel Fearns, Swetha Vijayakrishnan, David Bhella

## Abstract

Human respiratory syncytial virus (RSV) causes severe respiratory illness in children and the elderly. Treatments for RSV disease are however limited and efforts to produce an effective vaccine have so far been unsuccessful. Understanding RSV virion structure is an important prerequisite for developing interventions to treat or prevent infection but has been challenging because of the fragility of virions propagated in cell culture. Here we show, using cryogenic electron microscopy (cryoEM) and cryogenic electron tomography (cryoET) of RSV particles cultivated directly on transmission electron microscopy (TEM) grids, that there is extensive helical symmetry in RSV filamentous virions. We have calculated a 16 Å resolution three-dimensional reconstruction of the viral envelope, targeting the matrix protein (M) that forms an endoskeleton below the viral membrane. These data define a helical lattice of M proteins, showing how M is oriented relative to the viral envelope and that helical ordering of viral glycoproteins that stud the viral envelope is coordinated by the M layer. Moreover, the helically ordered viral glycoproteins in RSV filamentous virions cluster in pairs, which may have implications for the conformation of fusion (F) glycoprotein epitopes that are the principal target for vaccine and monoclonal antibody development. We also report the presence, in authentic virus infections, of N-RNA rings packaged within RSV filamentous virions. Overall, the structural data obtained provides molecular insight into the organization of the virion and the mechanism of its assembly.

## Introduction

Respiratory syncytial virus (RSV) causes acute lower respiratory tract disease in infants and the elderly. It is estimated that there are approximately 33 million RSV infections annually in children under the age of five years, leading to around 3 million hospital admissions and 60,000 in-hospital deaths. The great majority of these cases (~30 million) occur in low- and middle-income countries^1^. Although less common than seasonal influenza virus infection, RSV has been shown to be associated with higher mortality rates in hospitalised elderly patients^2,3^.

RSV is formally classified as Human orthopneumovirus within the *Orthopneumovirus* genus, *Pneumovirus* family and Mononegavirales order^4^. Closely related to RSV within the *Pneumoviridae*, in the *Metapneumovirus* genus is another notable pathogen, human metapneumovirus (HMPV) that also causes bronchiolitis and pneumonia in infants^5^. Thus, both ortho- and metapneumoviruses are clinically important respiratory pathogens.

RSV is a non-segmented negative sense RNA containing enveloped virus. The viral genome is 15.2 kb and has ten open reading frames encoding at least eleven gene products. The viral RNA is encapsidated by multiple copies of the viral encoded nucleocapsid protein (N) to form a left-handed helical ribonucleoprotein complex (or nucleocapsid - NC). This serves as the template for RNA synthesis by the RNA dependent RNA polymerase (RdRp)^6,7^, thought to occur in virus induced cytoplasmic organelles called inclusion bodies^8,9^. The RdRp comprises two proteins: the catalytic large (L) protein and the phosphoprotein (P) that mediates the interaction with the NC^10^. RNA synthesis is also modulated by the M2 gene products M2-1 and M2-2^11,12^. RSV encodes four envelope associated proteins, of which three are membrane proteins: the small hydrophobic protein (SH), fusion protein (F) and attachment protein (G). Underlying the envelope is the fourth envelope associated protein – the matrix protein (M), which coordinates virion assembly together with M2-1. M2-1 forms a second layer at the virion interior, under the M-layer, and associates with NCs^13,14^. High resolution structures for some of the envelope associated proteins of both RSV and HMPV have been determined by X-ray crystallography, including the matrix proteins^15–17^ the F glycoprotein^18,19^ and M2-1^20,21^.

At present, 39 vaccines and monoclonal antibodies (mAbs) targeting RSV are in development, the vast majority of which are directed at two conformations of the RSV F glycoprotein; pre-fusion and post-fusion, and their most neutralization-sensitive antigenic sites, Ø and V^22^. The RSV pre-fusion conformation of the F glycoprotein is the predominant form displayed at the virion surface and mediates viral entry by fusing viral and host cell membranes. The shapes of enveloped virions and the organization of their surface glycoproteins can impact virus biology by affecting fusion processes^23^, mucus penetration^24^ and complement activation^25^. A comprehensive understanding of RSV virion structure will therefore inform development of effective interventions to prevent RSV disease. Although the structures of RSV proteins and glycoproteins are becoming increasingly well understood, placing these structures in the context of the virion remains challenging, owing to the highly pleomorphic nature of virus particles purified from cell-culture^13,14^. Schematic diagrams mostly represent the RSV virion as a spherical particle, it has however, long been known to form predominantly filamentous virions at the assembly site^26^. We have previously shown that propagating filamentous enveloped viruses in cells grown directly on transmission electron microscopy grids leads to improved preservation for imaging by cryogenic electron microscopy (cryoEM)^27^. Here we employ this approach to image RSV filamentous virions by cryoEM and cryogenic electron tomography (cryoET). We show that although RSV virions are prone to loss of integrity, they are nonetheless highly ordered. Fourier analysis shows that filamentous particles exhibit helical symmetry. CryoET combined with sub-tomogram averaging led to the calculation of a three-dimensional density map of the viral matrix layer at 16 Å resolution. Fitting the X-ray structure of M shows a curved lattice of dimers that closely matches the arrangement seen in the (010) plane of the published X-ray structure (that is, the plane containing the *a* and *c* axes of the *C*2 unit cell). Our model orients the X-ray data relative to the virion filament axis and the viral envelope, providing a view of how M coordinates virion assembly. Our data also show that the matrix layer coordinates helical ordering of the viral glycoprotein spikes on the virion exterior. Finally, we show that glycoprotein spikes cluster in pairs on filamentous virions but can also pack in an alternate lattice in non-filamentous virions. Our data also demonstrate the presence of an abundance of ring-shaped assemblies, likely formed of the nucleocapsid protein N and RNA. Our findings concerning the structure of RSV provide a molecular understanding of virion organisation and assembly, information that is critically important in the development of effective vaccines or interventions to prevent or treat this major respiratory pathogen.

## Results

### CryoEM of filamentous RSV virions reveals helical order

To image RSV filamentous virions in as close to native conditions as possible we propagated virus in cells cultured directly on holey carbon support films. Over the course of our investigations several cell types were used – Vero (African green monkey kidney cells), U-2 OS (human osteosarcoma cells) and A549 (human adenocarcinoma cells). All were found to be suited to imaging filamentous virions. Imaging of filaments was optimal at 72 hours post-infection (fig 1A). Cryo-EM grids were therefore prepared by plunge-freezing at this time-point. Initial imaging of virus propagated in U2-OS and Vero cells in a JEOL 2200 FS cryomicroscope revealed well preserved filamentous virions, although it was clear that they are prone to physical disruption (fig S1A). Power spectra calculated from cropped sections of filamentous virions indicated a high degree of helical order evidenced by the presence of multiple layer-lines (fig S1B-C). To collect images better suited to Fourier analysis, RSV virions produced in U2-OS cells were imaged in a Thermo-Fisher Titan Krios 300 keV cryomicrocope, equipped with a Falcon II direct electron detector (fig 1B). The diameters of filaments extracted from these images varied considerably and spanned a range of 900 - 1600 Å. Fourier transforms (fig. 1C) showed patterns of layer lines consistent with helical symmetry. Indexing of the transforms using the program PyHI (personal communication Xuewu Zhang – University of Texas Southwestern Medical Center) suggested that the Bessel orders of the two principal maxima were very large, consistent with the tubes being constructed of many helical strands, probably of the order of 50. Helical objects such as these can be thought of as crystalline sheets that have been rolled up to form a tube^28^ and in a helix with this symmetry the units would lie on a lattice that can be specified by the two vectors indicated by the principal maxima, one of which had a length of ~86 Å aligned at ~85° to the helix axis and the second had a length of ~51 Å and aligned at ~49° to the helix axis. There was no density for the principal maximum for this second vector, the lattice was however established by using higher order reflections. Although the filtered image (Fig 1D) showed many areas where an indication of the underlying lattice of subunits could be discerned, the local lattices were only clear in limited areas, presumably a consequence of local distortions. This, together with the relatively weak intensity of the transforms and the large radii of the principal maxima frustrated the production of reliable 3-dimensional reconstructions using Fourier-Bessel methods. Such problems are often seen with helical tubes containing large numbers of strands^29^ and led us to explore whether cryo-ET could provide a structure of the underlying lattice.

**Figure 1.**
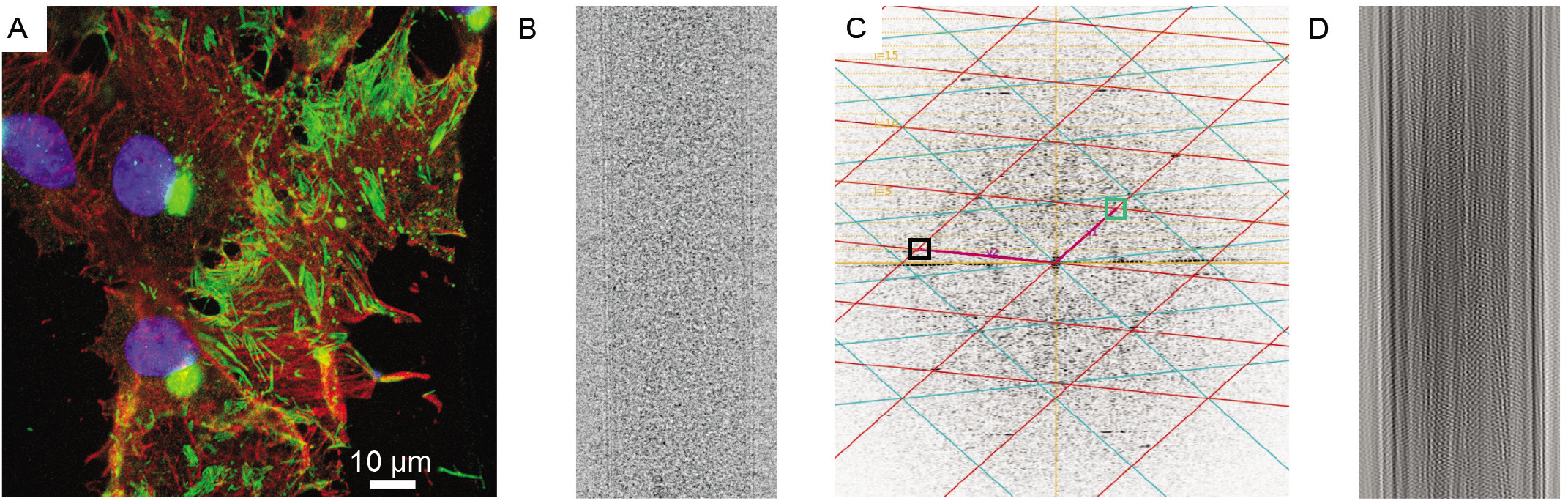
Respiratory syncytial virus forms filamentous virions that exhibit a high degree of helical order. (A) Immunofluorescence imaging of vero cells infected with RSV at 72 hours post-infection shows an abundance of filamentous particles. Virus particles were identified by labelling with an antibody that targeted the nucleocapsid protein (green). Phalloidin was used to localise actin (red) and DAPI reagent was used to stain cell nuclei (blue). (B) CryoEM image of a short section of RSV filament. (C) The power-spectrum for the filament shown in (B) is shown, Fourier-Bessel analysis was used to identify a putative helical lattice, the positions of principal maxima are indicated by boxes, however one was found to be missing (green box). The lattice was only able to be identified using higher-order reflections. (D) Masking and Fourier synthesis led to the reconstitution of a filtered image, showing that although a helical lattice is present, it is not sufficiently ordered to show features along the full length of the filament.

### Cryo-ET of RSV filamentous virions

To resolve the helical lattices underpinning RSV virion structure and assembly, we used cryogenic electron tomography to calculate three-dimensional reconstructions of individual virions. Fifty-eight tilt series were recorded on a Thermo-Fisher Titan Krios 300 keV cryomicroscope equipped with a Gatan K2 bioquantum energy filtered direct electron detector. Tilt-series’ of images were collected specifically targeting straight, undisrupted segments of filamentous virions. Low-magnification views of regions selected showed that although filamentous structure was well preserved by culturing virus on the TEM grid (fig 2A), many virions were to some extent disrupted, frequently having varicosities at points along their length or at one end. These data confirm that RSV virions are very fragile and easily lose their filamentous shape (fig 2B-C).

**Figure 2.**
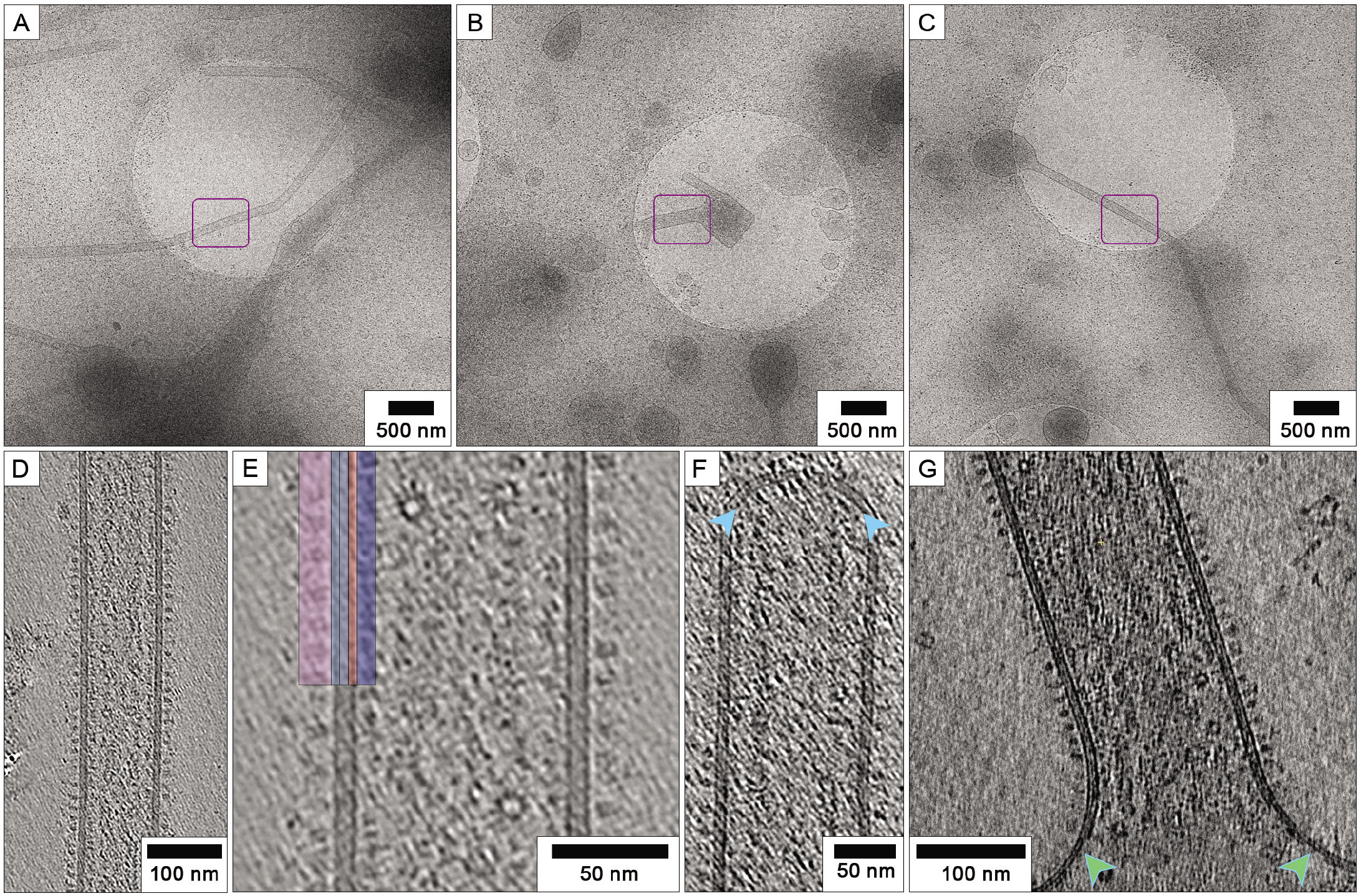
Cryoelectron tomography of RSV filaments reveals a complex envelope. (A) propagating RSV directly on the TEM grid led to improved preservation of filamentous morphology. (B,C) Despite the gentle preparation methods, many virions showed signs of disruption such as varicosities. The areas indicated by purple boxes were imaged to collect tilt-series’. The filament in panel A is shown in figure 3A, the filament in panel B is shown in figure 2G and the filament shown in panel C is shown in figure 3B. (D) a central slice through a denoised tomogram shows a complex viral envelope, shown at higher magnification in (E). Glycoprotein spikes are densely packed, giving a picket-fence like appearance when viewed from the side (highlighted in mauve). The lipid bilayer is highlighted in pale blue, the matrix layer in orange and the M2-1 layer in dark blue. (F) tilt-series’ were collected targeting straight regions of filamentous virions, however some tomograms captured the virion ends, which were found to be hemispherical but rather flattened. In some cases (as shown here) the matrix layer was intact and contiguous at the filament ends (blue arrows), while some virions showed an absence of M. (G) Loss of virion integrity was accompanied by discontinuities in the matrix layer (green arrows).

### A complex, ordered viral envelope

Tomographic reconstruction of filament sections reveal that they are well-ordered. As previously reported^13,14,30^, we find that the viral envelope is densely packed with viral glycoprotein spikes. Underlying the lipid bilayer is a contiguous density that we attribute to the matrix protein (M). This gives the viral envelope the appearance of being triple layered (fig 2D-E, movie S1 timepoint 0m 25s). Beneath the M-layer, is a second less sharply defined protein layer that has previously been attributed to the M2-1 protein^13^. Filament ends were typically hemispherical having a large radius of curvature, lending the caps a flattened appearance. Most tomograms of virion ends showed a contiguous matrix layer (fig 2F), although some were seen to have disrupted or missing matrix layers. Discontinuity in the matrix layer was more apparent in the varicose areas, indicating that loss of virion integrity is likely a consequence of detachment or mis-assembly of the matrix layer (fig 2G).

### Nucleocapsids appear as classical herringbone assemblies, loosely coiled helices and rings

The virion interior is densely packed with viral nucleocapsids, mainly having the characteristic herringbone morphology^31^ and suggesting that in common with several other mononegavirales, RSV virions are polyploid^32^ (fig 3A, movie S1 timepoint 1m 08s). Virions were also seen to be filled with an abundance of ring-like assemblies that ranged in diameter from 10 to 17nm (fig 3B). Upon close inspection of these structures, we found several rings that had radial spikes and morphology highly reminiscent of the decameric and undecameric rings produced by heterologous expression of the RSV N protein in insect cells or bacteria ^7,31,33,34^ (fig 3C movie S1 timepoint 1m 30s). In some cases, these ring-like features appeared to precess, when moving through successive sections of tomograms, whereas in others there was no obvious continuity of density (movie S1 timepoint 2m 20s). We hypothesise then that some of these ring-shaped assemblies are likely loosely coiled NC, but the absence of connecting density in successive tomogram sections and the presence of some isolated rings outside the virions (suspended in the vitreous ice layer - movie S1 timepoint 2m 05s) strongly suggest that many of these objects may indeed be N-RNA rings, perhaps being products of aborted genome replication.

**Figure 3.**
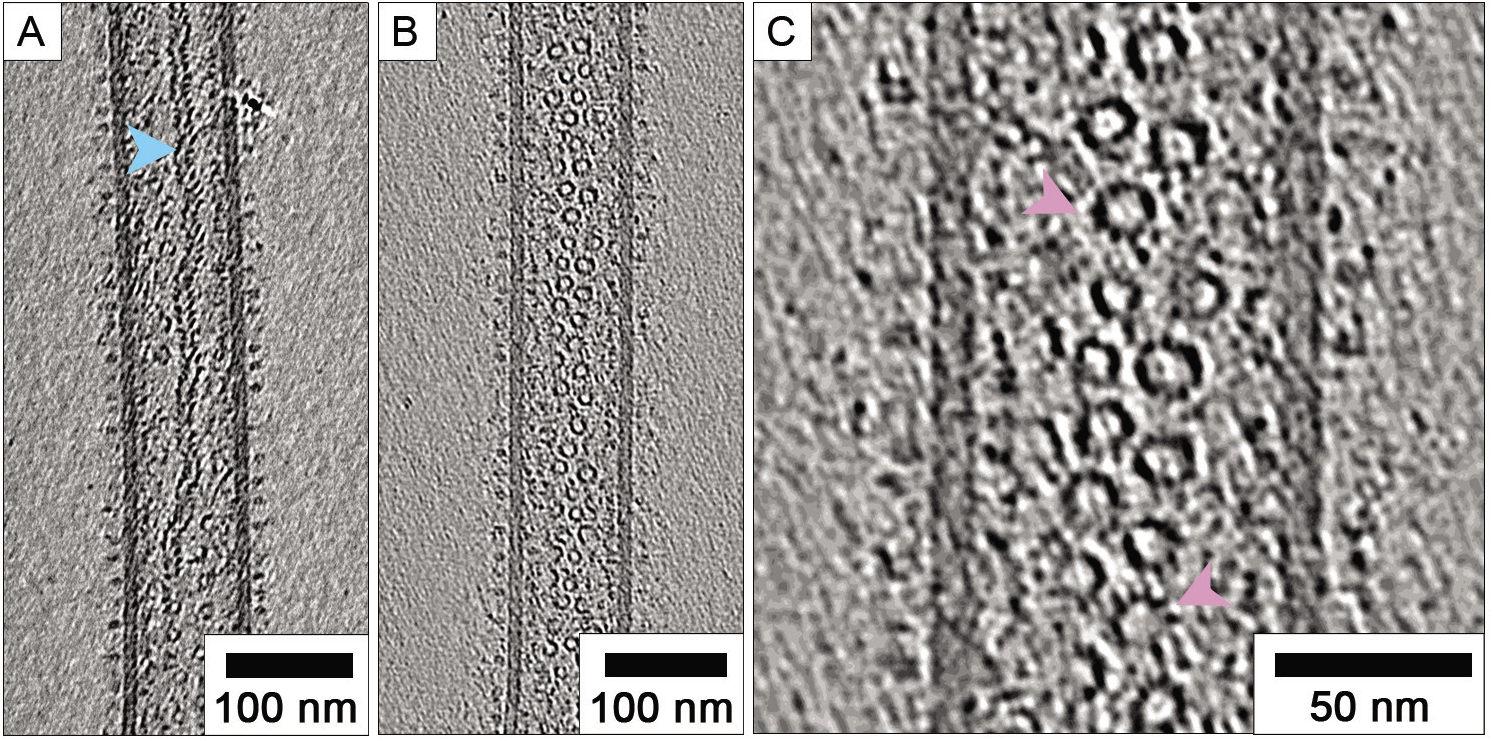
RSV virions package helical nucleocapsids and an abundance of N-RNA rings. (A) central slices through tomograms showed that virions contain multiple nucleocapsids, many having the characteristic herringbone morphology (blue arrow). (B) Many virions also contain large numbers of ring-shaped assemblies. (C) a close-up view of these structures reveals the presence of radial spokes (pink arrows). These structures are morphologically very similar to previously described decameric and undecameric rings produced by recombinant expression of RSV N.

### Helical ordering, clustering, and hexagonal arrays of glycoproteins

Previous studies have reported that RSV glycoprotein spikes are densely packed, giving a picket-fence appearance when viewed from the side (e.g. fig 2E) ^13,14,30^. A significant proportion of our tomograms showed helical ordering of glycoproteins, manifesting as striped arrays of density when viewed as slices taken through the glycoproteins at the top and bottom of virions (Fig 4A, movie S1 timepoint 2m 42s). Measurements of the spacings between the rows of glycoprotein spikes ranged between 89 and 208 Å, having both a mean and mode of 135 Å (SD=20.6, n=150). Some virions exhibited extensive and long-range ordering of the glycoproteins. Interestingly, the glycoprotein spikes were often observed to cluster in pairs (fig 4B). This was even more apparent in tomograms of virions that were only sparsely decorated with glycoproteins (fig 4C-F). Centre to centre measurements of the spacing of paired glycoprotein spikes gave a mean distance of 84 Å (SD=7.9, n=100).

**Figure 4.**
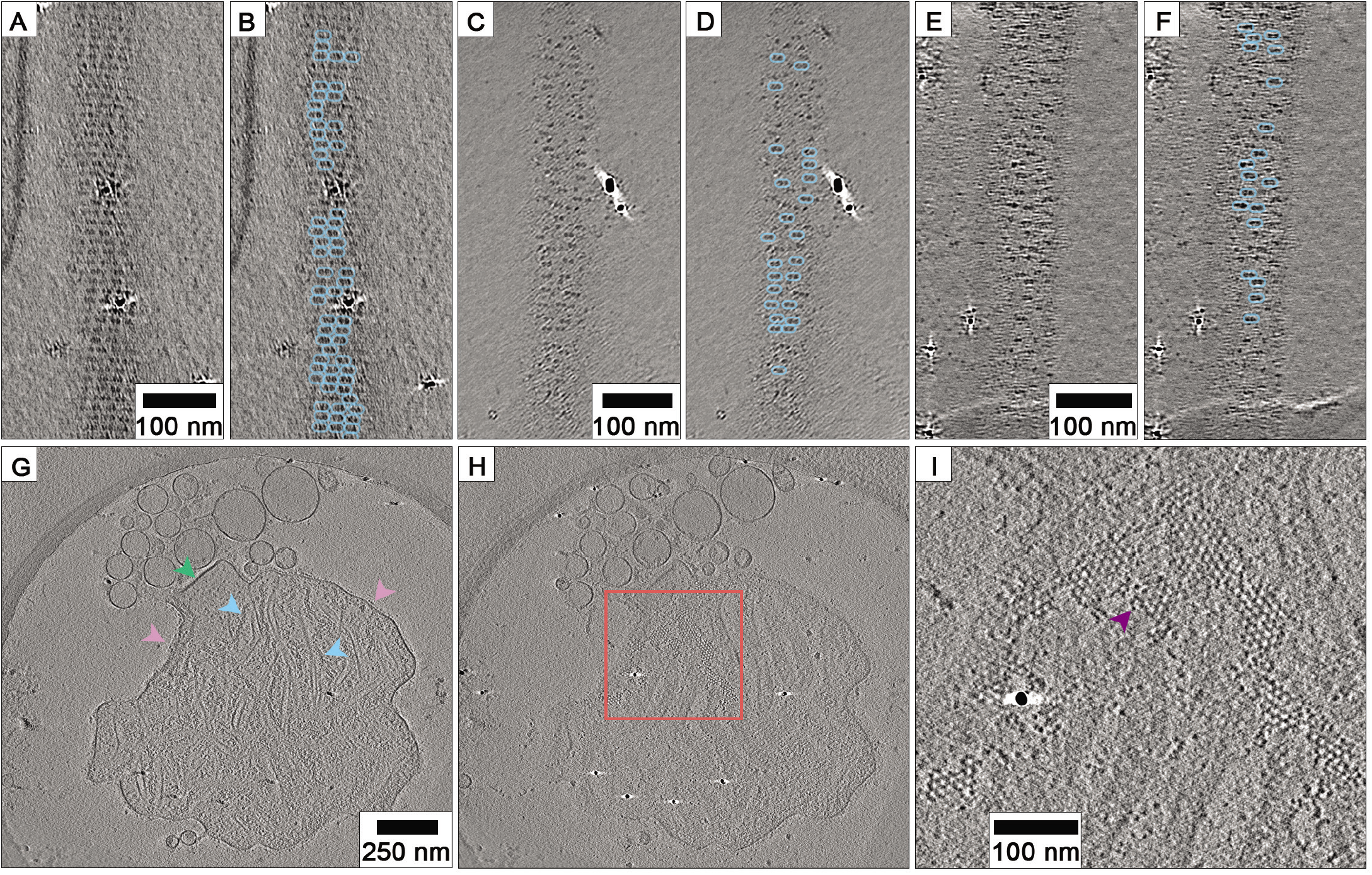
Helical ordering and clustering of RSV glycoprotein spikes. (A) a slice through the glycoprotein spikes of a filamentous RSV virion shows that they are helically ordered. Long range ordering of the glycoprotein spikes is evident, as is clustering of spikes in pairs (highlighted in (B)). (C-F) in virions that are sparsely decorated with glycoprotein spikes, clustering in pairs is even more apparent. (G) a central section through a tomogram of a pleomorphic RSV virion showing the presence of both herringbone nucleocapsids and N-RNA rings (blue arrows), flattened, thicker membranes that likely have a matrix-layer (green arrow), and tightly packed glycoprotein spikes (pink arrows). (H-I) a section through the envelope region of the same particle shows the presence of a honeycomb-like array of densities that may be an alternate arrangement of glycoprotein spikes, discrete densities are seen (purple arrow) that are likely individual glycoprotein spikes.

An earlier tomogram of RSV infected Vero cells, collected on a JEOL 2200 FS cryomicroscope showed a large membranous structure, that might be interpreted as a pleomorphic virion or cell debris. This object showed angular, thicker regions of membrane, consistent with the presence of a matrix layer (as also seen in fig 2B), picket-fence like arrangement of glycoprotein spikes, and contained both nucleocapsid and ring-shaped assemblies (fig 4G – movie S1 timepoint 3m 22s). This object also had patches of density packed to form a honeycomb lattice (fig 4H-I). The lattice appeared to be contiguous with the side-views of the glycoprotein spikes (movie S1 timepoint 3m 37s) and in places individual densities consistent with single glycoprotein spikes could be seen. Considering the clustering of RSV glycoprotein spikes in pairs described above, we hypothesise that this array is an alternate arrangement of glycoproteins. Making accurate measurements of the lattice was challenging as the densities were mostly not sharply resolved, nonetheless by measuring distances between vertices of hexagons we estimated the mean spacing between glycoprotein spikes to be 74 Å (SD=9.5, n=50).

### Sub-tomogram averaging of RSV envelope components – packing of matrix proteins

To better understand the way the RSV matrix protein drives virion assembly, we sought to calculate a three-dimensional reconstruction of the matrix layer by sub-tomogram averaging. Tiles of the viral envelope were extracted and aligned using Dynamo ^35^, yielding a 3D reconstruction of the matrix layer at 16 Å resolution (Fig S2). Viewed from the virion interior, the reconstruction reveals a regular array of M proteins (fig 5A-C, movie S2 timepoint 0m 7s). Viewed from the virion exterior the reconstruction has weak stripes of density that are visible at lower isosurface threshold levels (fig 5D-F, movie S2 timepoint 0m 32s). This feature is likely incoherent averaging of the viral glycoprotein spikes.

**Figure 5.**
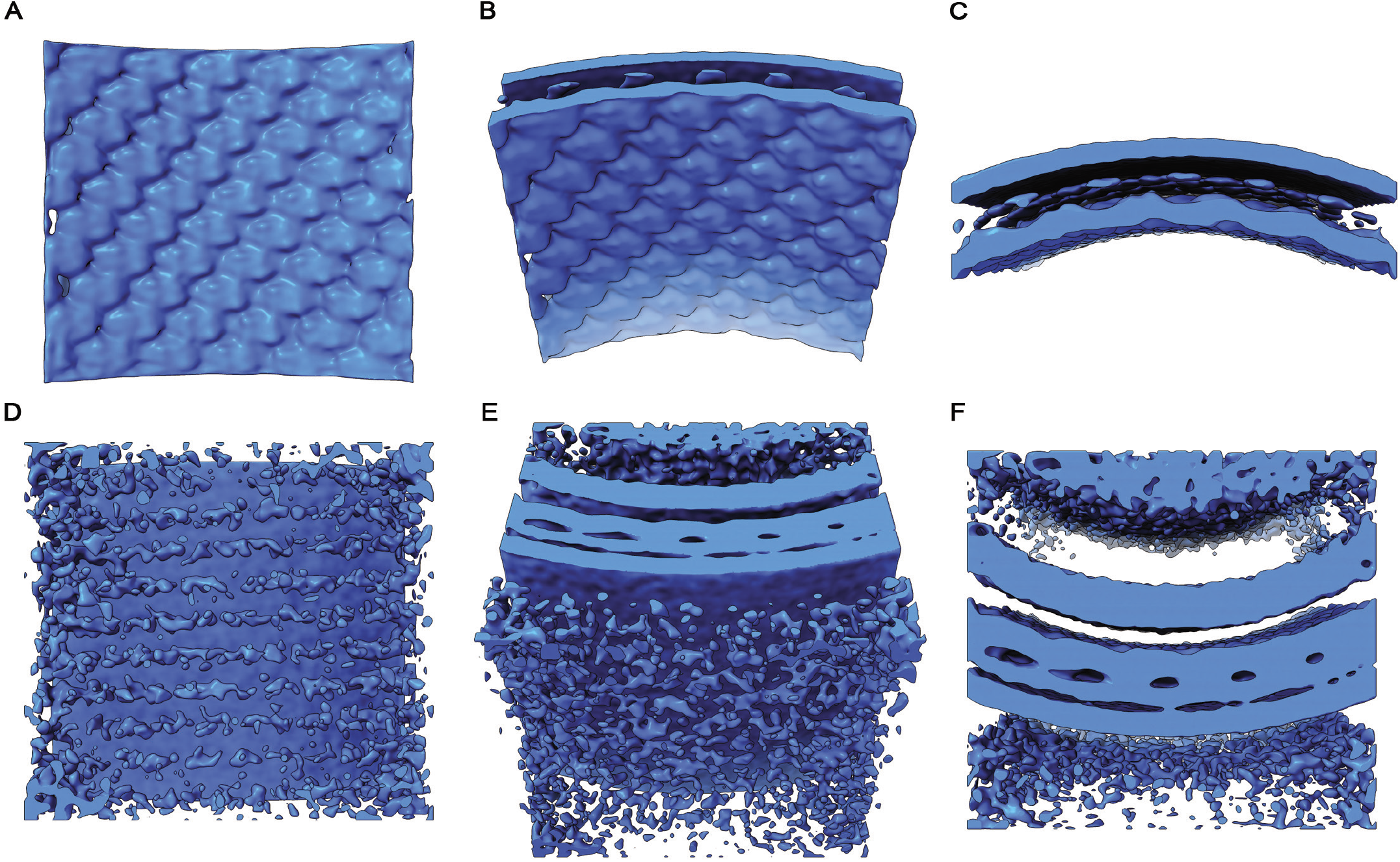
Subtomogram averaging of the RSV viral envelope reveals the structure of the matrix layer. (A) viewed from the filament interior, a regular array of density is seen in the envelope layer attributed to the matrix protein. (B) the average is tilted 45° towards the viewer. (C) the average is tilted a further 45°, giving a view of the lipid bilayer. The inner leaflet of the bilayer is only weakly resolved compared to the outer leaflet. (D) Viewed from the virion exterior at a lower isosurface threshold, stripes of weak, noisy density are seen. This is likely incoherent averaging of glycoprotein spike density. (E) tilting the reconstruction towards the viewer by 45° shows that the inner leaflet contacts the M-layer. (F) A fourth density layer is now visible, under the matrix. This density has been previously attributed to the RNA polymerase co-factor M2-1. Like the glycoprotein density at the exterior of the envelope, the M2-1 density is weak and incoherently averaged.

The X-ray structure and solution studies of the RSV M protein showed that it assembles into dimers^15^ (fig 6A-B, movie S2 timepoint 0m 46s). Interestingly in that study, the M dimers in the *C*2 crystal lattice were stacked into sheets (fig 6C, movie S2 timepoint 1m 10s). Docking the coordinates of the M-dimer (PDB 4V23) into our map generated a curved array that closely resembled the arrangement seen in the (*a,c*) plane of the X-ray structure (fig 6D-E, movie S2 timepoint 1m 54s). Measurements taken from this fitting experiment match closely those made in our Fourier-Bessel analysis (fig 6F). A low-pitch, right-handed helix follows the (1,0,-1) direction of the *C*2 unit cell of the X-ray structure, which is the (1,1) direction in the P2 sheet (shown in fig 6C). This had a spacing of 82 Å and an angle of 86.5° relative to the filament axis. The *a* and *c* axes in the *C*2 X-ray structure unit cell correspond to the long-pitch helical strands having rise and tilt measurements of 54 Å / 40° (left-handed) and 66 Å / 45° (right-handed) respectively.

**Figure 6.**
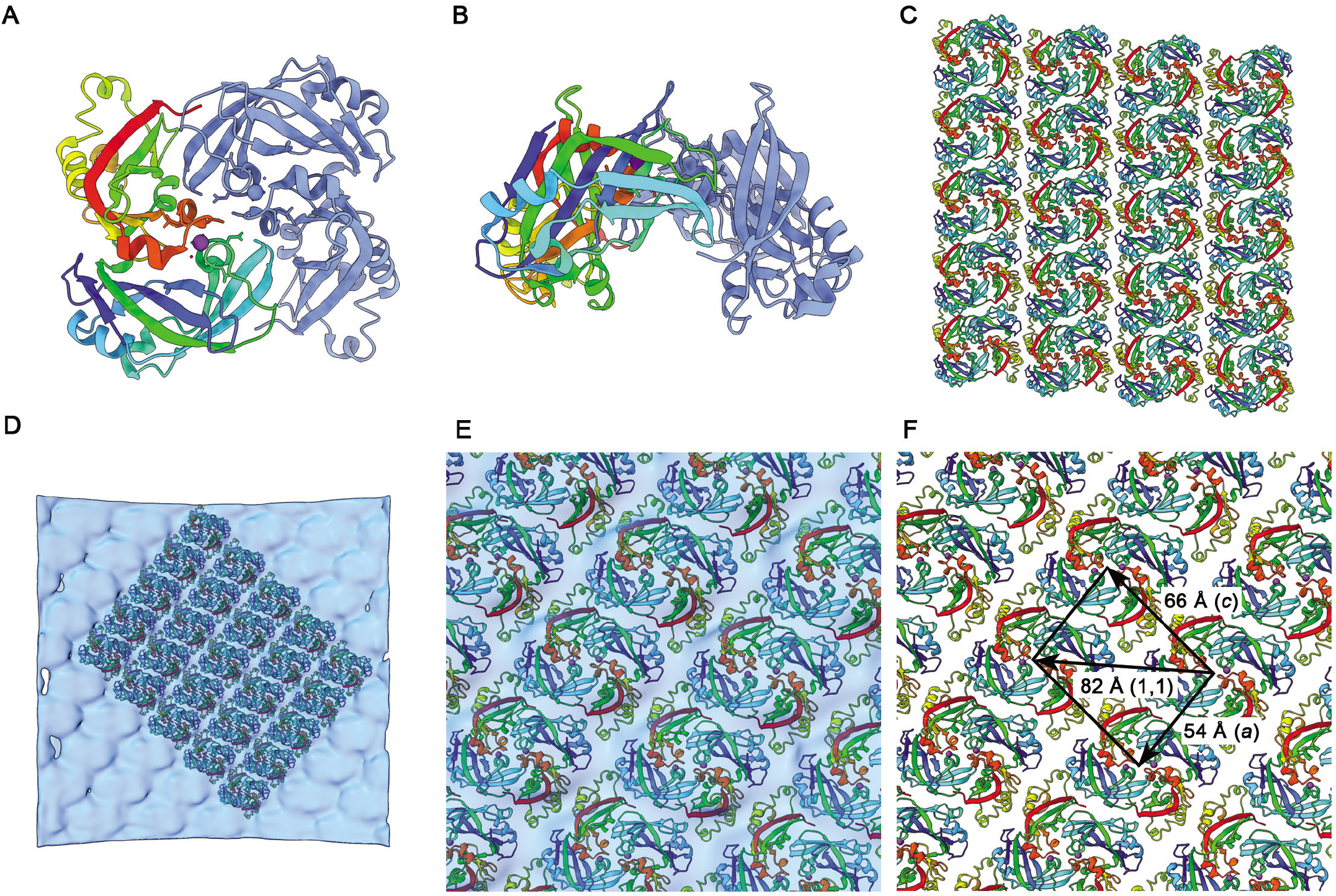
Docking the X-ray structure of M into the sub-tomogram average. (A-B) The published X-ray structure for RSV M shows that it forms a dimer (PDB accession number 4V23). (C) In the X-ray structure M dimers pack as stacked sheets in the (010) plane of the *C*2 crystals. (D) docking the M-dimer into the 3D reconstruction shows a curved lattice with similar packing *in virio* to the planar sheets seen in the crystal structure. (E) close up view of the docked M coordinates. (F) Measurements of the lattice spacings of M-dimers in the docked model of the matrix layer. A low-pitch helix corresponding to the (1,1) direction of the planar lattice (shown in panel C) has a spacing of 82 Å. Helix strands corresponding to the *a* and *c* axes in the *C*2 unit cell of the X-ray structure have spacings of 54 Å and 66 Å respectively.

Our fitted model clearly establishes the orientation of M-dimers relative to the viral envelope allowing the contact surface with the inner-leaflet to be identified (fig 7A). This surface is strongly positively charged (fig 7B-C movie S2 timepoint 2m 45s), whereas the interior surface of the matrix array presents stripes of negative charge (fig 7D-E, movie S2 timepoint 2m 35s) that may be important in coordinating the packaging of M2-1 and viral nucleocapsids. When considered in the context of the sub-tomogram average viewed at lower isosurface threshold, the stripes of density that we have assigned to the viral glycoprotein spikes align to the array of M-dimers (fig 7F). The spacing of these stripes of density does not however match those in the raw tomograms (45 Å versus 135 Å). These data suggest that the helix of glycoprotein spikes is coordinated by the matrix layer, albeit the array does not exactly follow the underlying lattice of M dimers. Rather, the process of aligning sub-tomograms to the lattice of the matrix layer has led to superposition of the more widely spaced array of glycoprotein spikes, leading to these densities appearing to be more closely packed in the sub-tomogram average than they actually are.

**Figure 7.**
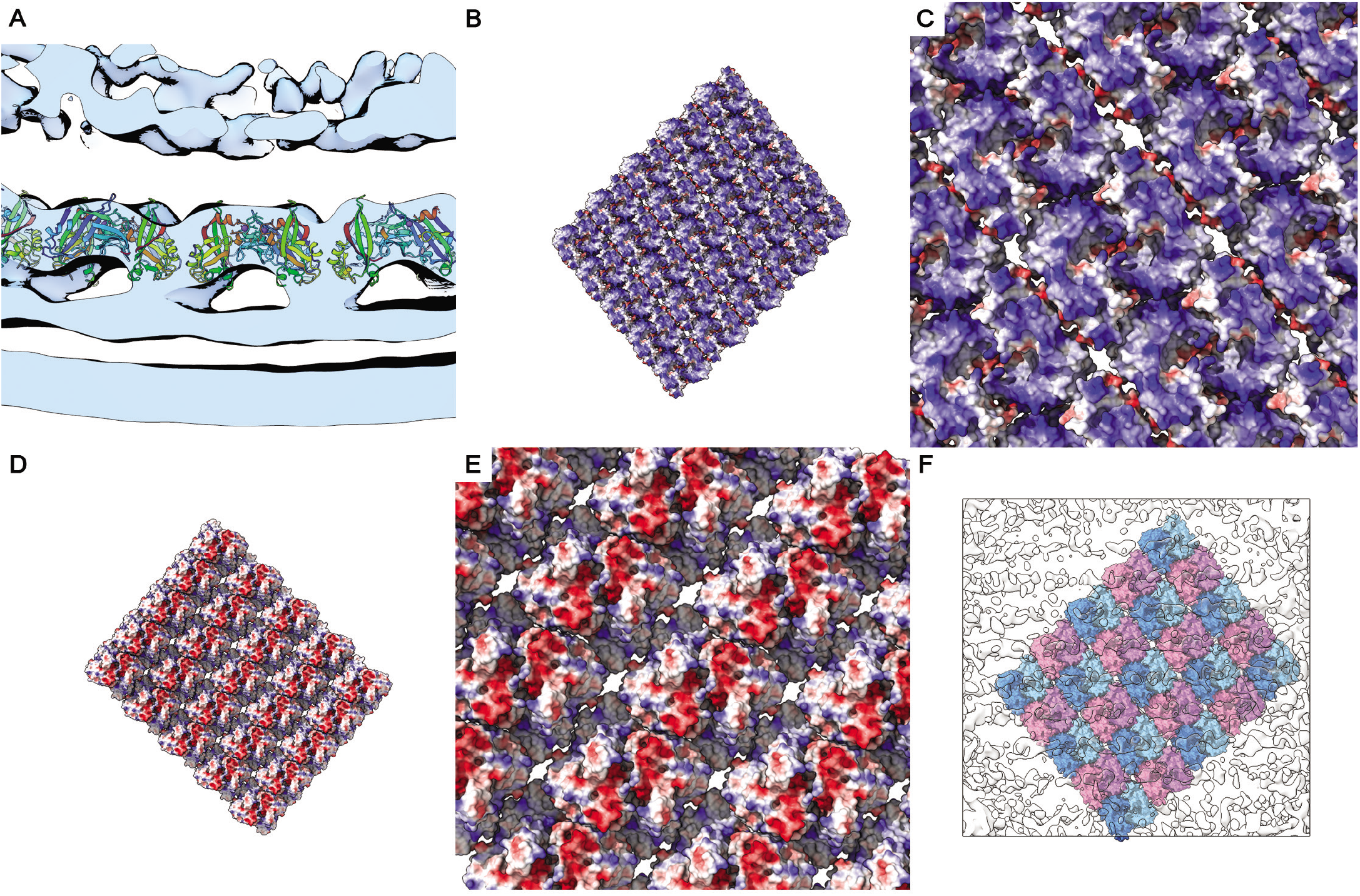
Orienting the matrix layer relative to the viral envelope and glycoproteins. (A) a section through the subtomogram average reconstruction with fitted coordinates for M shows the orientation of the M-dimer relative to the inner leaflet of the lipid bilayer. (B-C) a solvent excluded surface representation of the matrix layer is presented coloured to show the electrostatic potential. It shows that the surface facing the lipid bilayer is positively charged, while the surface facing the virion interior presents stripes of negative charge (D-E). (F) a view of the docked model placed within a transparent isosurface of the subtomogram average reconstruction and viewed at an isosurface threshold of 0 from the virion exterior shows that the noisy weak density that we attribute to the viral glycoprotein spikes aligns with the (1,1) helix (alternate strands are coloured pink and blue).

## Discussion

We have used cryogenic electron microscopy and tomography to study the structure of filamentous virions of respiratory syncytial virus propagated directly on the transmission electron microscopy support grid. This approach has allowed these large fragile structures to be imaged while minimising disruption caused by sample preparation. These data show that far from being a disordered pleomorphic virion comprising a membranous bag filled with nucleocapsid, RSV assembles highly ordered virions. Fourier analysis demonstrated that virions have extensive helical order. We were able to characterise the extent of this order by 3D imaging of virions using cryo-ET, combined with sub-tomogram averaging.

### Nucleocapsid morphology and packaging of N-RNA rings

Denoised tomograms of RSV filamentous virions revealed an abundance of nucleocapsids packaged within the virions. Many were seen to exhibit the classical herringbone morphology previously described for nucleocapsid-like particles produced by heterologous expression of the RSV N protein^6,31,34^. We also observed large numbers of ring-shaped assemblies. These rings bear a striking resemblance to N-RNA rings also previously described and although rings are produced following heterologous expression of N proteins from a variety of mononegavirales^31,36,37^ we believe that this is the first report of N-RNA rings being produced in authentic virus infections. It is not known what RNA species is associated with virion-associated N-RNA rings. Some of the most abundant RNA species in RSV-infected cells are short 21-25 nt RNAs generated from the leader (*le*) and trailer (*tr*) promoter regions^38–40^. Previous studies have shown that a fraction of these RNAs are nuclease resistant, indicating that they can be encapsidated^40^. Although it is well accepted that N-RNA rings would be expected to contain 70 nt of RNA encapsidated by 10 N protomers^6,7^, it has been shown that if purified recombinant N protein is incubated with RNAs as short as 14 nt, it can form N-RNA rings that are of similar dimensions as those reconstituted with longer RNAs^41^. This suggests either that RNA is only required to nucleate the encapsidation event and that subsequent N-N interactions allow formation of a 10-protomer ring that is not necessarily entirely RNA-bound, and/or that short N-RNA complexes can associate together to form 10 N-protomer rings containing multiple small RNA oligonucleotides. Thus the ~20-25 nt RNAs found in abundance in RSV infected cells could potentially be incorporated into N-RNA rings and packaged into virions. Whether N-RNA rings fulfil a functional role in newly infected cells, such as nucleating liquid-liquid phase separated biomolecular condensates to serve as transcription/ replication centers^42,43^ or inhibiting cellular stress responses^44^ remains to be determined.

### Packing of M-dimers in the matrix layer and variations in filament radii

Fourier analysis of cryoEM images of filamentous virions showed the presence of helical ordering in these particles. The highly variable radii of filamentous virions together with the weak intensity of layer lines and missing principal maxima in the Fourier transforms frustrated our efforts to compute a 3D reconstruction using Fourier-Bessel methods. Nonetheless, these data encouraged us to continue investigating the structure of these important virions using tomography and sub-tomogram averaging. This led to the calculation of a 3D reconstruction of the viral envelope that showed how M-dimers pack together to form the matrix layer, unequivocally orienting the dimers relative to the inner-leaflet of the lipid bilayer and orienting the M lattice relative to the virion’s filament axis. Docking the X-ray structure to our cryo-EM 3D reconstruction showed that the curved lattice of M-dimers closely matches the planar arrays previously seen in the M crystals. In figure 6F we show measurements of vectors along the lattice, a right-handed (1,1) helix with a spacing of 82 Å between subunits and a left-handed (*a*) helix with a spacing of 54 Å. The axial rise per subunit for each of these helices is approximately 5 Å and 45 Å respectively. If the matrix tube was to polymerise as a simple 1-start, right-handed (1,1) helix then it would comprise 8 helical strands (along the *a* or *c* directions) and have a radius of ~100 angstroms. This is considerably smaller than the measured radii of the matrix layers in our viral filaments, which ranged between 370 and 660 Å (fig S3). Thus, we expect the matrix layers of filamentous virions to exhibit more complex geometries assembling as *n*-start (1,1) helices and incorporating different numbers of helical strands (likely between ~30 and ~50). The matrix layer can be thought of as a sheet corresponding to the (*a,c*) plane of the *C*2 crystal lattice, that has been rolled into a tube. Because the number of helical strands in a filament is large, only small distortions would need to be introduced to bend the sheet to form a cylinder. Indeed, distortions of the M dimer interface that were postulated to foster curvature have been observed^15^. Incorporating different numbers of helical strands in the matrix layer would accommodate the considerable variation in filament diameters that are observed. Although each different number of strands would be associated with different helical parameters, the underlying M lattice would remain the same.

### Helical ordering and clustering of glycoprotein spikes – implications for virion assembly, entry, and the design of interventions to prevent RSV disease

Our tomograms showed several virions that exhibited clear helical ordering of glycoproteins, features confirmed by sub-tomogram averaging. Measurements of the spacing between successive turns of these helices gave a mean value of 135 Å. The spacing between strands of the low-rise right-handed (1,1) helix of M dimers that coincides with the stripes of density attributed to the glycoprotein spikes in our sub-tomogram average was 45 Å however. This, together with the weak intensity of this feature in the reconstruction (usually a consequence of low-occupancy and/or incoherent averaging), indicates that the helical ordering of the glycoprotein spikes is coordinated by, but not congruent with the underlying lattice of matrix proteins. The most likely explanation of this anomaly is that the helical array of glycoprotein spikes is coordinated by every third (1,1) helical strand (and less frequently by every second or fourth strand). Whether this phenomenon is simply a consequence of steric collision preventing association of a glycoprotein spike with every M-dimer, or an allosteric mechanism whereby binding of a glycoprotein spike to one matrix dimer favours binding to successive dimers on the same strand, or prevents binding on adjacent strands, remains to be determined. It may even be the case that the underlying interaction between M and M2-1 regulates the interaction between M and the cytoplasmic tails of the glycoproteins.

Intriguingly we have found that glycoprotein spikes tend to cluster in pairs. This finding may have considerable implications for our understanding of both viral attachment and entry processes, as well as epitope presentation and vaccine design. Further investigation targeting the structure of paired glycoprotein spikes will be necessary to establish whether they comprise F-F, G-G or F-G clusters, and whether clustering is coordinated by M, or an intrinsic property of the glycoprotein spikes. The spacing of glycoprotein spikes in doublets on filamentous virions was measured at 84 Å, whereas the spacing of M-dimers along the low-pitch helix measured 82 Å. Thus, doublet formation may be coordinated by M packing, the slightly larger spacing being a consequence of the measurements being made at a higher radius. Sub-tomogram averaging using masks to target both the glycoprotein spikes *and* M from tomograms of virions that have very well-ordered glycoproteins may therefore provide a more detailed view of the interaction between M-dimers and the cytoplasmic tails of glycoproteins and confirm whether M coordinates glycoprotein spike doublet formation as well as helical packing.

In addition to helical ordering of glycoprotein spikes with doublet formation, we also observed formation of an alternate lattice of glycoprotein spikes, seen on a large pleomorphic virion. Measurements of this array indicated closer packing of spikes (74 Å), although the analysis was subject to greater ambiguity, owing to the less sharply resolved density. It would be interesting to establish whether this mode of packing reflects ordering of glycoprotein spikes at the plasma membrane and whether it too is coordinated by M. It has previously been shown that M is critical for the formation of filaments but not nucleation of budding sites^45^. Association of M with detergent-resistant lipid microdomains (thought to be the sites of virion assembly) has been shown to depend on glycoprotein expression^46,47^. Confocal microscopy showed the presence of M protein in inclusion bodies and at the cytoplasmic side of the host-cell plasma membrane^47^. One possible interpretation of the honeycomb-like packing of glycoprotein spikes may be that they (and possibly also M) have the potential to pack in a fullerene-like manner at the plasma membrane, fostering the formation of hemispherical caps. Filament elongation might then be driven by assembly of the helical M-dimer lattice, polymerising along the low-pitch helix and engaging the cytoplasmic tails of F and G to form the membrane-enclosed helically ordered virions we have observed. Further studies employing cryo-ET of RSV budding sites may define more precisely the ordering of envelope associated proteins and glycoproteins at the plasma membrane and thereby inform our understanding of virion morphogenesis.

### Summary

In the present study, we have used a range of cryogenic electron microscopy and image analysis approaches to characterise the structures of RSV filamentous virions. In so doing we have shown that RSV packages large quantities of N-RNA rings as well as full-length genome containing nucleocapsids. We have provided a detailed description of the viral envelope, in which we discerned the packing of M-dimers, describing how they are oriented relative to the inner-leaflet of the viral envelope and how they are ordered to form a helical array. Furthermore, we show that the helical packing of M-dimers coordinates helical ordering of viral glycoprotein spikes and that the spikes have a propensity to form doublets, a feature that may also be coordinated by M. Finally, we have described an alternate packing of glycoprotein spikes that may be important for virion morphogenesis.

Our findings indicate that future structural analyses have the potential to provide detailed insights into glycoprotein-matrix interactions that drive virion morphogenesis, information that may lead to interventions to treat RSV disease. Moreover, our discovery of extensive ordering of viral glycoproteins has implications for our understanding of the mechanisms of virus attachment and entry and may inform the design of more effective antigens for improved vaccines to prevent RSV disease.

## Methods

### Confocal microscopy

Vero cells were grown in Dulbecco’s modified eagle medium (DMEM, Gibco, Life Technologies, UK) supplemented with 10% foetal bovine serum (FBS; Gibco, Life Technologies) at 37°C in an atmosphere containing 5% CO2. Monolayers grown to 75% confluency on glass cover slips were infected with Human Respiratory Syncytial Virus (strain A2 - RSV) at a multiplicity of infection (MOI) of 0.5 for 1 hour and then washed, replacing the media with DMEM containing 2% FBS. At 24-, 48- and 72-hour time-points, cells were fixed in 4% formaldehyde and 2.5% Triton X-100 in PBSA, a blocking step of incubation with sheep serum for 1 hour was followed by immunostaining for N using a mouse monoclonal antibody (αN009 ^48^) and detected using a sheep anti-mouse FITC conjugate (Sigma, UK). Phalloidin Alexa Fluor 568 (Invitrogen, UK) was used to stain actin. Cells were mounted using ProLong Antifade plus DAPI reagent (Invitrogen, UK). Imaging was carried out with the Zeiss LSM710 laser scanning confocal microscope.

### Propagation of RSV on cryoEM grids

U2-OS or A549 cells were seeded on to cryoEM grids as follows. Freshly glow-discharged finder gold quantifoil grids (200 mesh R2/2 – Quantifoil MicroTools GmbH, Germany) were sterilized in 70% EtOH and placed in a glass-bottomed MatTek dish (MatTek Corporation, MA, USA). 200 μl of laminin (50μg ml^−1^) was added and incubated overnight. Grids were then washed in DMEM. 10^5^ cells were added in 2ml of DMEM supplemented with 10% FBS and incubated at 37°C in an atmosphere containing 5% CO2 overnight before infection. Cells were infected with RSV at a MOI of 1 and incubated for a further 72 hours before being prepared for cryoEM. Grids were frozen for cryoEM by plunge freezing. Briefly, 3μl of a suspension of 5nm colloidal gold beads were pipetted onto each grid (BBI Solutions, United Kingdom). Grids were then transferred to a Vitrobot Mk IV (Thermo-Fisher Scientific), blotted for four seconds and then immediately plunged into a bath of liquid ethane.

### Imaging – projection images

RSV virions were initially imaged on a JEOL 2200 FS equipped with a Gatan 914 side-entry cryostage and a Gatan Ultrascan 4000 CCD detector, operated at 200 keV. Energy filtered images were recorded with a slit-width of 10 eV at 50k× magnification, corresponding to a pixel size of 2.2Å. To improve the quality of images collected we sought access to a 300 keV cryomicroscope equipped with a direct-detection camera. Further cryo-EM was therefore performed at the MRC – Laboratory of Molecular Biology on a Thermo-Fisher Scientific Titan Krios microscope equipped with a Falcon II detector. The microscope was operated at a nominal magnification of 29k× giving a calibrated pixel size of 2.84Å at the specimen scale. Micrographs were recorded as movies, comprising 70 individual frames and with a dose per frame of 1 electron/Å^2^ and at 18 frames per second.

### Imaging - tomography

Initial tomography experiments were performed on a JEOL 2200 FS, as noted above. Continuous tilt-series were collected, using SerialEM, from −60° to +60° at 2° intervals and at a nominal magnification of 20k×, corresponding to an unbinned pixel size of 5.32 Å at the specimen scale. Improved tomography data, suitable for sub-tomogram averaging were then collected at the UK electron bioimaging centre at Diamond Light Source (eBIC) on a Thermo-Fisher Scientific Titan Krios microscope equipped with a Gatan BioQuantum K2 energy filtered direct detection camera. Dose-symmetric tilt-series collection was performed using SerialEM ^49,50^. Energy-filtered images were collected with a slit-width of 20eV and an applied defocus of between −2 and −4.5 μm. One second exposures were recorded with an electron exposure of 2 electrons/Å^2^, partitioned over five movie frames. A total of 41 images were recorded per tilt-series at 3° intervals from −60° to +60°, thus a total exposure of 82 electrons/Å^2^ was applied per tilt-series. Tilt-series were recorded at a nominal column magnification of 81k× corresponding to a calibrated pixel size of 1.79 Å at the specimen scale.

### Fourier analysis

Preliminary data collected on the JEOL 2200 FS were processed to determine the presence of helical layer lines using SPIDER^51^. Briefly, overlapping sections of a single filamentous virion were extracted. The log of the power spectrum for each image section was calculated. The power spectra were then summed. Higher-quality data collected at 300 keV and using a direct electron detection camera were visualised using Ximdisp ^52^. Indexing was performed using PyHI (personal communication Prof. Xuewu Zhang UT Southwestern medical centre). Fourier synthesis was used to produce a filtered image of the helical components by assigning crystallographic indices to pairs of reflections in the Fourier transform. The image was reconstituted by masking and inverse Fourier transformation using trmask, a program within the MRC image processing software suite^53^.

### Tomogram calculation

Individual movies were corrected for drift and beam induced motion using MotionCor2 ^54^. Motion corrected images were then compiled into an angle ordered stack file using a perl script to extract the correct files and tilt angles from the SerialEM metadata files (mdoc), creating the final tilt-series using the IMOD command newstack ^55^. Tilt-series alignment and reconstruction by weighted back projection was then accomplished using the IMOD package. Defocus estimation was performed using CTFFIND4 ^56^. For visualisation and interpretation tomograms were binned by a factor of four and then denoised using the denoise3d option in Topaz ^57^.

### Sub-tomogram averaging

Tomograms were recalculated with a SIRT-like filter (50 iterations) and 8× binning to assist in filament axis definition and particle picking in Dynamo ^35,58^. A catalogue was created with 10 tomograms containing 11 filamentous RSV virions. Sub-tomogram/particle coordinates were defined along the filaments using the ‘crop on rings along path’ model in Dynamo. From the 10 tomograms, 30,088 sub-tomograms were extracted from the oversampled coordinates in a box size of 32 voxels, oriented normal to the filament/viral envelope. An initial average was calculated and smeared along the filament major axis to produce a featureless reference. More accurate sub-tomogram positions were calculated by running an alignment project with the featureless reference and allowing shifts only in the z direction to locate the precise position of the viral envelope.

Sub-tomograms were aligned to a smeared and low pass filtered reference while allowing angular and rotational searches of 15° with 5° sampling. Shifts of 4, 4 and 1 pixels were permitted in X, Y and Z directions, respectively. Initial alignments were performed with a wide saddle shaped mask (to exclude density from the viral envelope and M2-1 layers) followed by a tighter mask. Upon completion of the alignment projects, the sub-tomograms were re-centred in the extraction boxes based on the improved coordinates and extracted from tomograms with a reduced level of binning i.e. 4×, 2× and 1× binning. 15 alignment iterations were performed at each binning level. The averaging and alignment protocols were repeated for each reduced level of binning with an increasing box size until unbinned averages were attained (in a 256^3^ box). For the final alignment step, sub-tomograms were extracted from tomograms calculated using weighted back-projection (i.e., omitting the SIRT-like filter). Half-maps were generated and compared using the Relion post-processing and local-resolution tests ^59^. A FSC cut off of 0.5 was adopted to measure the resolution of the map for the M-layer using a saddle shaped mask. Averages were visualised and X-ray structures were fitted in UCSF Chimera and ChimeraX ^60,61^.

## Acknowledgements

The authors wish to thank Drs. Greg McMullan and Kutti R Vinothkumar for support with data collection at the LMB. We thank Diamond Light Source for access to the Cryo-EM facilities at the UK national electron bio-imaging centre (eBIC), proposal EM16637-5, funded by the Wellcome Trust, MRC and BBSRC. We are indebted to Daniel Clare for his expert assistance during dose-symmetric tomography data collection at eBIC.

This work was supported by the United Kingdom Medical Research Council (MC_UU_12014/7 to DB; MR/M000451/ for DB, RF, MJC, JS and SEB; MC-A025-5PL41 for MS). RF, DB and AMB were supported by the United States of America National Institutes of Health (1R01AI113321), BJP and HJ were supported by a Wellcome doctoral training programme (102463/Z/13/Z and 099786/Z/12/Z respectively)

The MRC-University of Glasgow Centre for Virus Research uses the CRediT taxonomy of author contributions. MJC – formal analysis, writing – review, JMS – formal analysis, methodology, writing – original draft, JH – methodology, writing – review, AMB – investigation, JS – investigation, SEB – investigation, MS – formal analysis, writing – review, BJP – investigation, writing – original draft, HJ – investigation, GZ – methodology, writing - review RF – conceptualisation, funding acquisition, writing – original draft, SV – investigation, methodology, supervision, DB – conceptualisation, methodology, investigation, formal analysis, funding acquisition, supervision, visualisation, writing – original draft.

## Data deposition

The sub-tomogram average described in this paper and a representative tomogram have been deposited in the electron microscopy data bank – accession number EMD *****

## Notes

### Competing Interest Statement

The authors have declared no competing interest.

