## Supplemental figures for "Helical Ordering of Envelope Associated Proteins and Glycoproteins in Respiratory Syncytial Virus Filamentous Virions"

**Supplemental information**

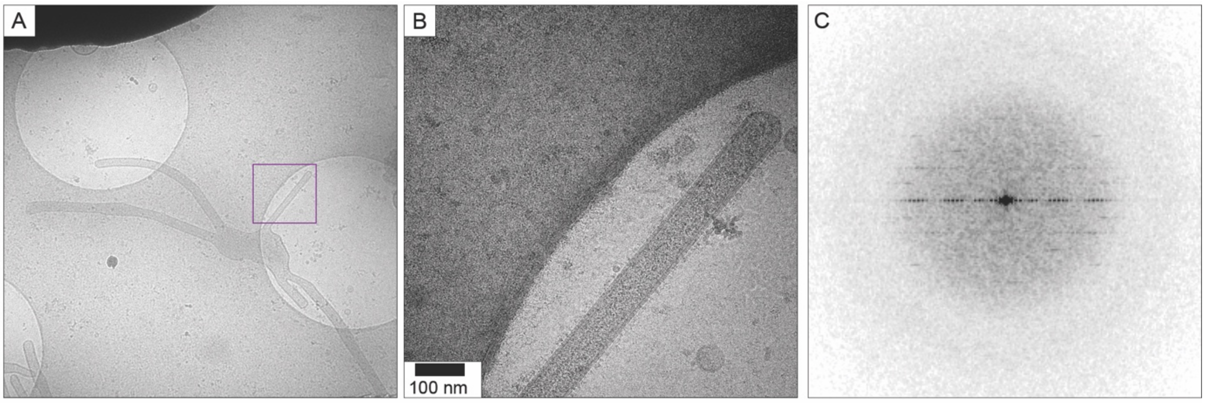


**Fig S1** *Preliminary imaging and analysis of RSV filamentous virions.* Imaging of virions propagated directly on the cryo-EM grid led to preservation of filamentous morphology. Virions were nonetheless prone to loss of filament integrity and formation of pleomorphic particles (A). Imaging (B) and Fourier analysis (C) of filamentous sections (boxed area in A) showed that these regions were highly ordered, as demonstrated by the presence of multiple layer-lines in the Fourier transform (C).

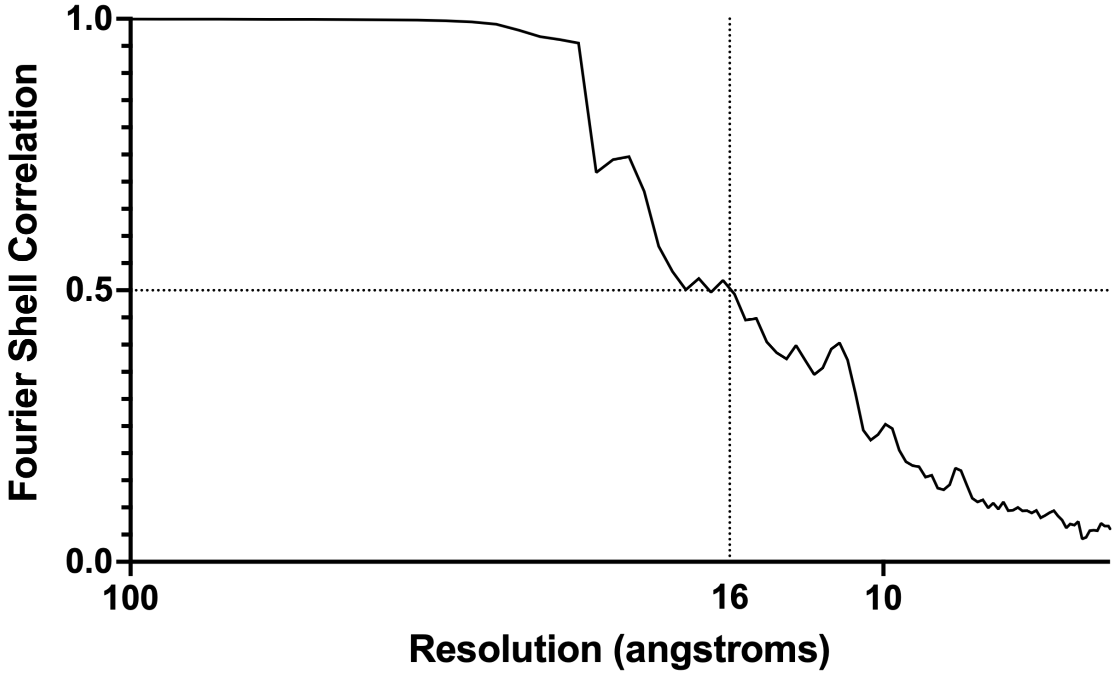


**Fig S2** *Resolution measurement for the sub-tomogram average of the RSV viral envelope.* Fourier shell correlation analysis showed that sub-tomogram averaging, focussed on the matrix array, achieved a resolution of 16 Å. Resolution assessment used a cut-off of 0.5, as the gold-standard protocol was not used during sub-tomogram alignment.

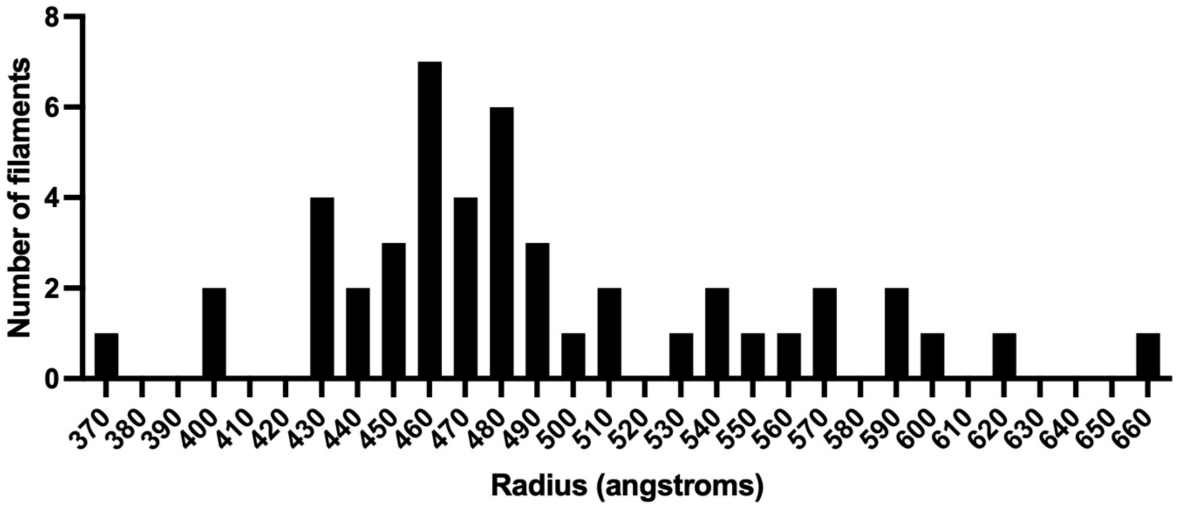


**Fig S3** *Radius measurements for the matrix layers in tomograms of RSV virions.* Filamentous virions show a considerable variation in radius. M-layer radii ranged between 370 and 660 Å and appear to show a multi-modal distribution.
